# Adaptive variation in avian eggshell structure and gas conductance across elevational gradients?

**DOI:** 10.1101/2022.11.07.515494

**Authors:** David Ocampo, Carlos Daniel Cadena, Gustavo A. Londoño

## Abstract

Many tropical species have restricted elevational distributions, which are potentially bounded by constraints imposed by physical conditions on physiological processes. Although some studies have examined variation in the physiology of adult birds with respect to elevation, little attention has been paid to the structure and function of eggshells, which mediate gas exchange between the embryo and the environment. At high elevations, dry air is expected to increase water loss from the egg; selection to avoid desiccation might therefore be expected to favor reduced gas conductance by means of increased eggshell thickness or reduced pore size. We used gas diffusion experiments and scanning electron microscopy to examine water vapor conductance rates and eggshell structures in 197 bird species distributed along an elevational gradient in the Andes. As predicted, water vapor conductance across the eggshell declined in a narrow range with elevation among all species and among species within families, but not among individuals within species. Variation among species in eggshell conductance was lower at high-elevation sites, potentially indicating greater constraints at such sites. Structural changes in eggshells with respect to elevation varied among taxonomic families of birds, suggesting potentially different adaptive responses to common selective pressures in terms of eggshell thickness and pore density, and size. We suggest that considering functional and structural traits of eggshells, which influence embryo development, may help one to better understand the elevational distributions of species and to forecast their responses to global climate change.

## Introduction

Species distributions along environmental gradients are constrained by their responses to biotic interactions and abiotic conditions (*e*.*g*. temperature, humidity). The role that such factors play in establishing the elevational distributions of tropical montane species has been increasingly studied, partly owing to an interest in forecasting species responses to climatic change (Sheldon et al., 2011; García-Robledo et al., 2016; Slatyer & Schoville, 2016; Zuloaga & Kerr, 2016; Freeman et al., 2018; Linck et al., 2021; Bota-Sierra et al., 2022). Because of their narrow elevational ranges and often abrupt replacement of closely related species along mountain slopes, tropical birds have been a model system for the study of elevational range limits (Terborgh, 1971; Diamond, 1973; Sheldon et al., 2011; Jankowski et al., 2013; Londoño et al., 2015; Freeman et al., 2019; 2022). Aside from biotic interactions, physiological constraints on adult individuals (*e*.*g*. heat and water loss, energetics, respiration) likely set elevational range limits in tropical birds (Janzen, 1967; Ghalambor et al., 2006; Cadena & Loiselle, 2007; White et al., 2007; McCracken et al., 2009; Storz et al., 2010; Jankowski et al., 2013; Londoño et al., 2015; 2017; Freeman, 2016). A seldom considered hypothesis is that physiological requirements of embryos may also restrict the ability of birds to breed over wide elevational ranges (Jankowski et al., 2013) because the partial pressures of oxygen, humidity, and temperature, which vary considerably with elevation, are key factors affecting embryo development (Carey et al., 1982; Deeming, 2002; 2011).

The exchange of oxygen and water vapor in bird eggs occurs by diffusion across the shell through microscopic pores (Tullett & Board, 1977). Thus, the evolution of avian eggshell structure involves a trade-off between oxygen uptake and water loss, and desiccation of the embryo, both of which are influenced by eggshell thickness and by the density and size of pores (Ar et al., 1974; Vleck et al., 1979). The eggshell structure has been studied in a few model species (Rahn et al., 1979), particularly with respect to egg size, but little is known about how eggshells vary among bird species across environmental gradients (Sotherland et al., 1980; Board, 1982; Carey et al., 1989). Likewise, although considerable work exists examining factors affecting gas exchange across eggshells (Sotherland et al., 1980; Carey et al., 1983), comparative analyses aimed at understanding the influence of environmental conditions on variation among species in eggshell function remain scarce (Vleck et al., 1983; Portugal et al., 2014; Attard & Portugal, 2021). More generally, there are few comparative studies of eggs involving multiple species aside from analyses focused on variation in egg shape (Stoddard et al., 2017; Duursma et al., 2018) and eggshell wettability and thickness (Attard et al., 2021; Attard & Portugal, 2022).

Studies on temperate-zone species have shown that the eggshell structure of birds may vary with temperature, humidity, and barometric pressure. Specifically, eggs of bird species that inhabit dry and cold high-elevation environments reduced eggshell pore density or pore size and increased eggshell thickness, which reduced the gas diffusion across the egg when compared to eggs of lowland species (Rahn et al., 1979; Rahn & Ar, 1980; Sotherland et al., 1980; Carey, 1980*a*; Rahn et al., 1982; Rahn & Paganelli, 1990). In contrast, in a population of a passerine species that recently colonized a high-humidity environment, eggs were larger, eggshells thicker, and pore density lower than in populations from drier habitats (Stein & Badyaev, 2011). These results are opposite to the pattern one would expect if gas exchange were the main driver of variation in eggshell traits; alternatively, the observed variation in eggs might reflect the elevated risk of trans-shell bacterial infection in highly humid conditions (Cook et al., 2005).

What little we know about eggshell structure in tropical birds has been generalized from studies on a limited number of taxa. Among ducks and grebes, for example, the eggshells of high-elevation species have lower pore density and lower water vapor conductance than those of closely related lowland species of similar size (Carey et al., 1982; 1983; Carey et al., 1990). In this study, we use phylogenetic comparative methods applied to measures of water vapor conductance and data obtained using scanning electron microscopy to examine patterns of variation among and within species in eggshell function and structure across elevational gradients in a variety of Neotropical landbirds.

Atmospheric pressure and the water vapor pressure of the atmosphere decrease with elevation, such that eggs are expected to lose water more rapidly at higher altitudes, all else being equal (Romanoff, 1930; Wangensteen et al., 1974). Accordingly, we tested the prediction that to retain water over the incubation period, eggshells of birds breeding at higher altitudes should exhibit lower rates of water vapor diffusion (Sotherland et al., 1980; Rahn et al., 1982). This could be accomplished by producing eggs with thicker shells (*i*.*e*. greater pore length), smaller pore diameter, or fewer pores relative to birds from the lowlands (Wangensteen et al., 1971; Board, 1982). We also asked whether eggshell function and structure may vary adaptatively within species due to mechanisms similar to those acting across species by assessing whether similar patterns of variation in water vapor diffusion rate and eggshell structure occur along elevational gradients in analyses at different taxonomic levels (*i*.*e*. among families, among species within families, among individuals within species).

## Methods

We worked at three biological stations along an elevational gradient in the Andes of Manu National Park, Cusco Department, Peru (Londoño et al., 2015): Pantiacolla (380 – 500 m elevation; 12°390’ S, 71°130’ W), San Pedro (1300 – 1600 m; 13°030’ S, 71°320’ W), and Wayquecha (2550 – 3200 m; 13°100’ S, 71°350’ W). We also sampled three sites in Colombia: Remedios in the mid-Magdalena Valley, Antioquia (500 m; 6°70’ N, 74°37’ W); Estación Biológica ICESI in Parque Nacional Natural Farallones de Cali (2200 – 2500 m; 03°26’ N; 76°39’ W); and the Cerro de Montezuma, Parque Nacional Natural Tatamá, Risaralda (1000 – 2600 m; 5°23’ N, 76°08’ W). In total, we studied eggs from 197 species of birds. We tested the assumption that habitats were drier at higher elevations, which increases the gas diffusion potential (Romanoff, 1930), by placing humidity/temperature data–loggers (HOBO pro V2 est temp/RH) along the elevational gradient in Peru (Low: Pantiacolla, Medium: San Pedro, and High: Wayquecha), during two breeding seasons (Jun-Dec: 2012–2013) (Supplementary data 1). The average ± SD ambient temperature per station was: low elevation = 22.5 ± 2.9 °C, mid-elevation = 17.6 ± 2.9 °C, high elevation = 10.7 ± 3.6 °C (Supplementary data -Figure S1b).

### Gas exchange experiments

We examined variation in gas diffusion rates across the eggshell in 141 eggs (one egg per nesting event) of 108 species (56 species from Peru and 52 species from Colombia; Table S1) by measuring daily water loss under known conditions of temperature, humidity, and barometric pressure (Ar et al., 1974). In contrast to earlier studies using eggshell fragments (Portugal et al., 2010; Maurer et al., 2011), we used entire eggs, which may more closely estimate gas flux under natural conditions given that shell thickness and pore density or size may vary along different regions of the egg (Rokitka & Rahn, 1987) and because water balance of entire eggs also involves effects of the inner membrane (Paganelli et al., 1973). Because avian eggs lose mass during incubation due to evaporation (Romanoff & Romanoff, 1949; Deeming, 2002), the rate of mass loss can be used to determine the permeability of the eggshell to gases (*i*.*e*. conductance; *sensu* Ar et al., 1974) if ambient water vapor pressure is known. We weighed recently laid eggs (<5 days after laying) collected from nests with known age that were abandoned or partially predated, each day for 7–9 days on a MyWeight balance accurate to 0.01 g. Most (ca. 80% from monitored nests) of the eggs were fresh (1-3 days after laying). For the other 20%, we were not able to confirm the laying date, but we are certain they did not have a developed embryo; only blood vessels were observed in these eggs (Fierro-Calderón et al., 2021). Eggs were placed in desiccators with silica gel (SiO_2_) to maintain humidity close to zero. Although the overall temperature was not strictly controlled given logistical difficulties in remote field sites, desiccators were kept at relatively constant temperatures at each field station (average ± SD; low elevation = 22.9 ± 2.9 °C, mid-elevation = 17.7 ± 3.5 °C, high elevation = 12.6 ± 2.4 °C) which do not allow avian embryos to develop (Funk & Biellier, 1944).

Water vapor conductance (GH_2_O mg * day^-1^ * torr^-1^; Ar et al., 1974) was determined as a rate of the water loss per day (24 hrs), corrected to a standard barometric pressure of 760 torrs because the rate of diffusion is inversely proportional to total pressure (Paganelli et al., 1971). The small reduction of vapor tension caused by the solutes of the egg contents can be neglected because the standard barometric pressure between the inside of the egg and outside the shell was close to zero (Paganelli et al., 1971). We did not detect temporal variation during the incubation period in water vapor conductance of the eggshells on the linear regression analyses performed (R^2^ > 0.8) for all the species.

Because each of the species was studied at only one of the field stations (*i*.*e*., they did not occur at different elevations), we calculated mean conductance for species where N>1, maximum of 3 eggs per species from different nests. Because gas conductance across eggshells may be affected markedly by the nest environment (Portugal et al., 2014), we sought to minimize the effect of nest types on our assessment of variation with elevation by sampling species using ground, cup, dome, and cavity nests in similar proportions across elevations (χ2 = 8.34, *df* = 6, *P* = 0.215). We performed generalized linear models (GLMs) to assess the effect of egg weight (mass), elevation (station), nest type, and latitude (*i*.*e*., country) on conductance, and we ranked models using Akaike’s Information Criterion accounting for small samples sizes (AICc). In addition, our analyses focusing on individual families (see below) control for nest type, as closely related species typically have similar nest structures.

### Characterizing eggshell structure

We examined eggshell structure of eggs from birds in Peru with scanning electron microscopy (SEM). We photographed eggshell fragments (ca. 3 × 3 mm) using a Philips XL30 SEM at the Smithsonian National Museum of Natural History (Washington DC, USA) and measured eggshell structure variables (see below) in 147 species (one egg per species) belonging to 30 families distributed along the elevational gradient (Table S2). Because we were interested in examining the eggshell surface under natural conditions, eggshells were not cleared nor stained (Fig. 1; Fecheyr-Lippens et al., 2015).

**Figure 1.**
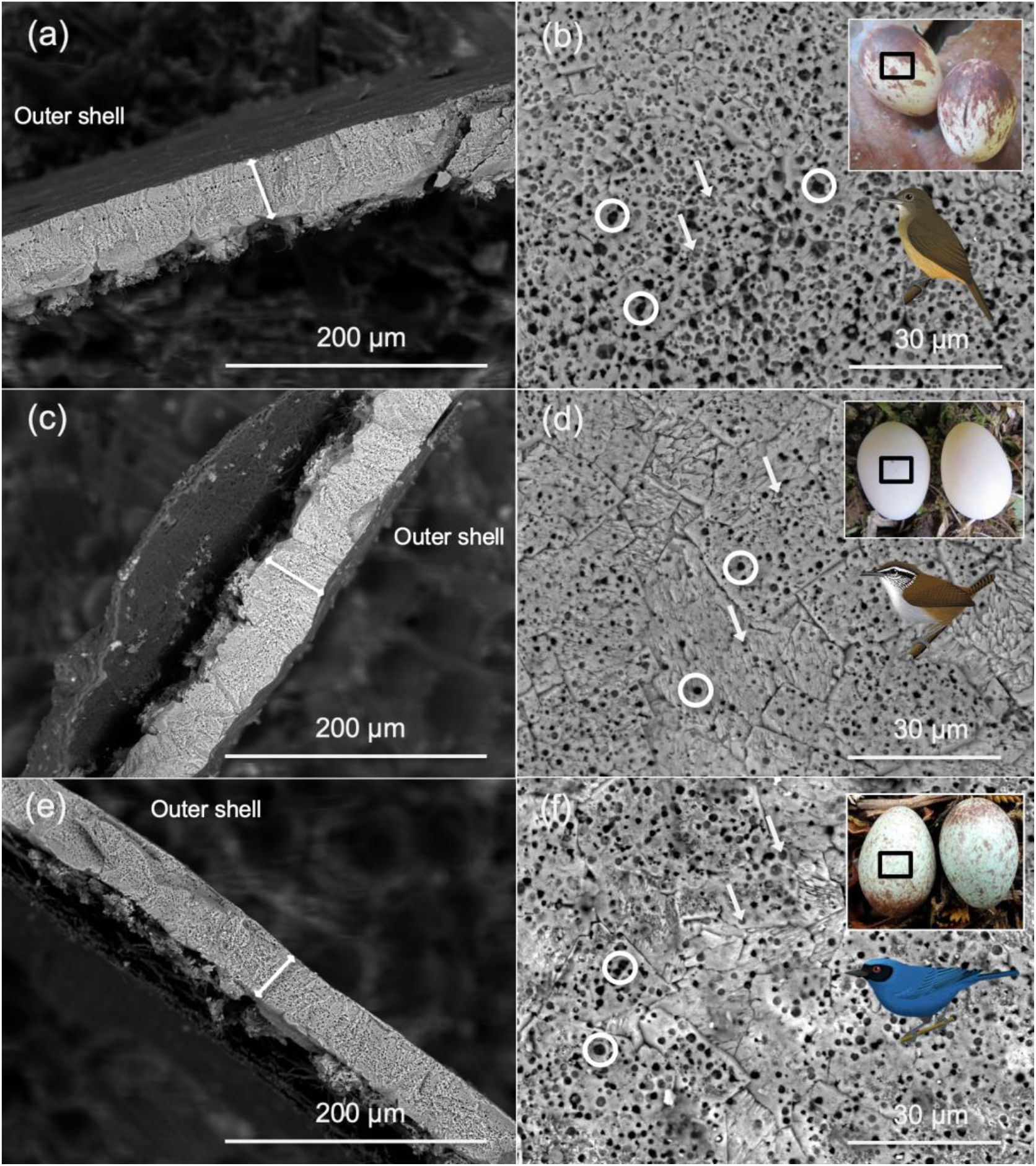
SEM images of the cross-section (a, c, e) and the eggshell surface and eggs species photo/illustration (b, d, f) of Dusky-throated Antshrike (*Thamnomanes ardesiacus*, a and b), Gray-breasted Wood-Wren (*Henicorhina leucophrys*, c, and d) and Masked Flowerpiercer (*Diglossa cyanea*, e, and f). White double arrows in a, c and e show the thickness measurement taken from the eggshell; white circles in b, d, and f point to exposed pores counted, and white arrows in occluded holes. Photos by DO and Birds illustrations by Fernando Ayerbe-Quiñones.

#### Shell thickness

We took one photograph of the shell cross-section (Fig. 1a, c, d) in 129 species. Using ImageJ (Rasband, 2008), we took three separate linear measurements of shell thickness and used the mean value as our estimate of eggshell thickness. To avoid biases related to variation in embryo development, we did not measure the mammillary layer of the eggshell because for some groups its thickness decreases with embryonic development (Karlsson & Lilja, 2008; Österström & Lilja, 2011).

#### Pores

To estimate pore density and size, we photographed the surface of three eggshell fragments from the equatorial region of each egg (Fig. 1b, d, f) in 116 species (one per species) for which pores were visible (in several non-passerines, pores were covered by cuticle structures on the surface of the eggshell, Mikhailo (1997); D’Alba et al. (2016)). We developed an image-processing algorithm (Correa & Hernández et al., unpublished) in MATLAB (2013a, The MathWorks, Inc., Natick, Massachusetts, United States), which automatically recognizes and quantifies pores in SEM images, and measures their mean sectional area (*i*.*e*., size in μm^2^). We assumed that functional pores were the dark holes, that contrast with the gray holes that appear occluded (Rahn et al., 1979; Board & Scott, 1980; Vieco-Galvez et al., 2021; *see* Figure 1). With these data, we calculated pore density and pore size (Table S2).

### Data analysis

All variables were log-transformed prior to analysis to account for nonlinear scaling. We accounted for size-related effects by regressing GH_2_O (water vapor conductance) against egg mass and eggshell characteristics (thickness, pore density and mean pore size) against egg volume (mm^3^), and then using residual values (when regressions were significant) in subsequent analyses. Egg volume was calculated based on the length (*L*) and width (*W*) of each egg as V = (0.6057 − 0.0018*W*)\**LW*^2^ (Narushin, 2005). We used mass or volume in different analyses depending on the availability of data for each eggshell trait (*i*.*e*. we did not have mass data for all eggs). To account for egg size in analyses of GH_2_O we used mass because it was widely used in past studies. Because mass and volume are highly correlated, however, using one or the other should not generate any directional bias in the results for our comparative purposes (Paganelli, et al., 1974).

We assessed whether conductance and eggshell traits exhibited phylogenetic signal (Pagel, 1999; Freckleton et al., 2002) and tested whether patterns of evolution in these variables are consistent with a process involving selection. This allowed us to describe the mode of trait evolution (*see below*) and to select the most appropriate model for testing for variation in traits with elevation. To build the phylogeny for these analyses, we randomly selected 500 phylogenetic trees from BirdTree (Jetz et al., 2012) using the Hackett et al., (2008) backbone phylogeny and used the *fitContinuous* function in the GEIGER library in R (Harmon et al., 2007) to obtain maximum-likelihood values and parameter estimates for four models of evolution: Brownian motion (Felsenstein, 1973), change concentrated at the tips (lambda; Pagel, 1999), early-burst (EB; Harmon et al., 2010), and constrained random walk with a central tendency (Ornstein–Uhlenbeck process with a single optimum; Butler & King, 2004). Based on the Akaike Information Criterion, the Ornstein–Uhlenbeck model best fit our data and was used for all comparative analyses (Table S3).

We compared residual (*i*.*e*., size-independent) eggshell traits among field station elevations using phylogenetic ANOVA as implemented in the ‘caper’ package for R (Orme, 2013). We performed the analysis combining the data collected in Peru and Colombia, using three elevation categories (low –ca. 340m-850m–, mid –ca. 1200m-2000m–, and high –ca. 2500 m-3000m–). Similar results were obtained when data from Peru and Colombia were analyzed separately. Additionally, we ran analyses separately for families with >3 species included in our sample: Emberizidae (5 species), Icteridae (4 species), Thraupidae (15 species), Turdidae (7 species), and Tyrannidae and allies (15 species) to ask whether different lineages may be responding to common pressures in different ways.

To test for variation in eggshell traits consistent with the hypothesis of adaptation to conditions varying with elevation, we examined the relationship between residual mass-independent variables and elevation with phylogenetic general linear models (pgls) implemented in the ‘phytools’ package for R (Revell, 2010; 2012). In addition to examining patterns across the complete data set, we evaluated whether similar patterns existed within different taxonomic groups. We examined relationships between elevation and eggshell variables separately in families having data for >2 species at different elevations: Caprimulgidae (5 species), Emberizidae (4 species), Furnariidae (8 species), Grallariidae (3 species), Thraupidae (12 species), Trochilidae (13 species), Turdidae (genus *Turdus*, 4 species), and Tyrannidae and allies (19 species). Additionally, we tested for cross-elevation trait variation among individuals within one species of Passerellidae (Black-faced brush finch, *Atlapetes melanolaemus*) and one species of Tyrannidae (Cinnamon Flycatcher, *Pyrrhomyias cinnamomeus*) for which we found nests over a relatively broad elevational range (*n* = 13 nests, range 2300 – 3000 m for *A. melanolaemus*; *n* = 19 nests, range 1300 – 3000 m for *P. cinnamomeus*).

## Results

According to our predictions, high elevations presented significantly lower relative humidity (*F*_2,94368_ = 7062, *P* < 0.001, Supplementary data–Figure S1a). During the breeding season, nesting birds at high elevations encounter dry environmental conditions, where 15% of the data was below a threshold of 80% of relative humidity. In contrast, only 3.1% and 3.2% of the data recorded at medium and low elevations, respectively, were below 80% of relative humidity.

## Conductance

We found a low phylogenetic signal (Pagel’s λ = 0.38) in eggshell conductance (GH_2_O), suggesting this is an evolutionarily labile trait. Our best model (ΔAICc ≤ 2) was the one including elevation and egg weight (EW) as predictor variables (GH2O ∼ 1 + Elevation + EW; AICc= 834.2457, weights =0.6461). Larger eggs lose more water compared to smaller eggs, and higher elevation eggs lose less water than medium and low elevations (*slope* = 0.56, *r*^2^ = 0.27, *P* < 0.001; Fig. 2a)., supporting our predictions related to the importance of elevation.

**Figure 2.**
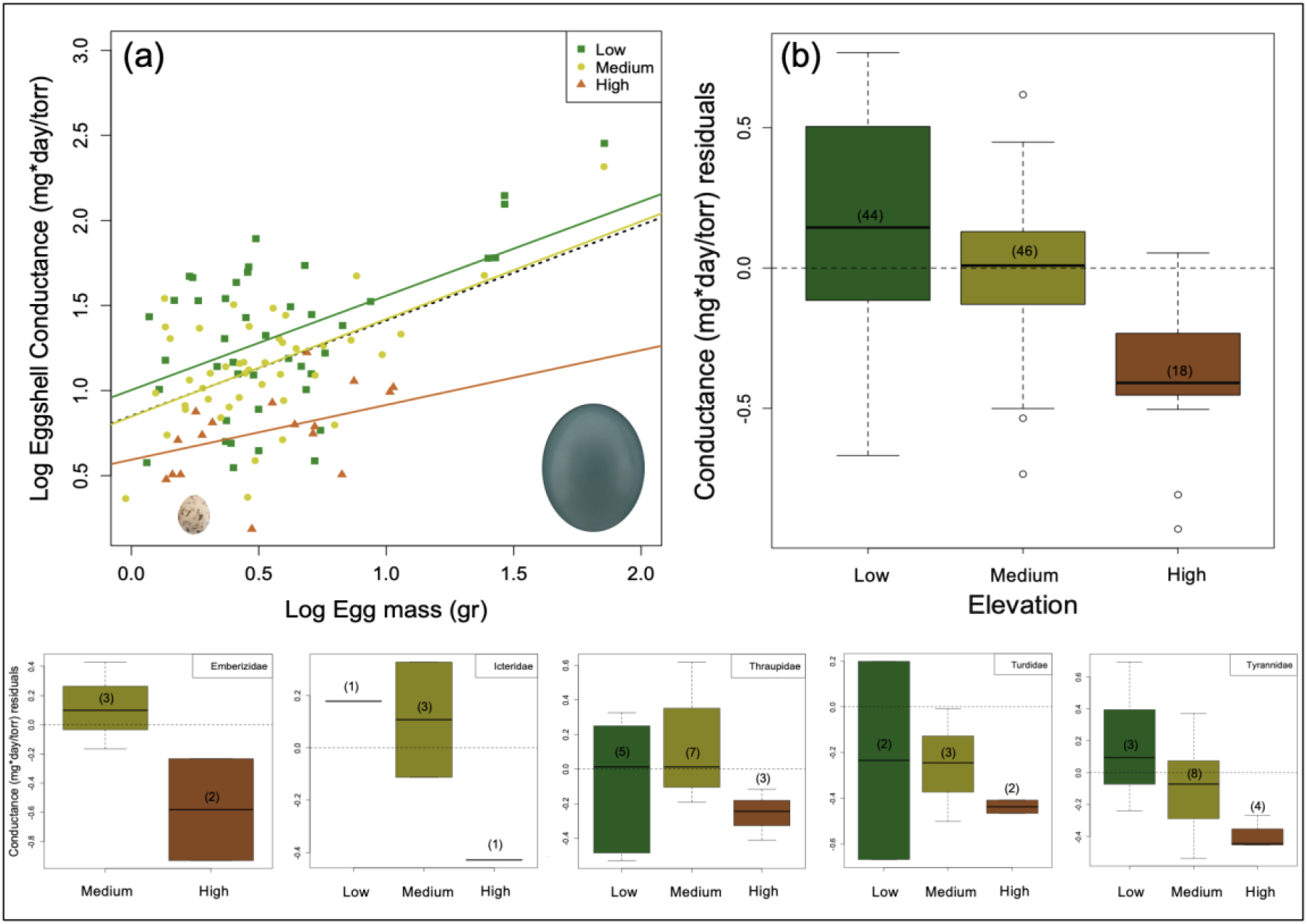
(a) Positive correlation (*F*_2,106_ = 29.77, *P* < 0.001) between egg mass and eggshell conductance across 103 species of birds over parallel elevation gradients in Peru and Colombia. Dotted lines indicate the general trend per elevation. (b) Residual conductance of species occupying three different elevational ranges (Phylogenetic ANOVA; *F*_2, 106_ = 4.46, *P* < 0.001; lowland *n* = 44; medium elevation *n* = 43; and highland *n* = 16); note that lowland species show wide conductance values with a mean above the expected given the allometric relationship. Highland species exhibit less variation and the conductance is overall lower than expected for the size of their eggs. Below, residual conductance at the three different elevations are shown separately by family; note the same pattern of low conductance at high elevation. Sample sizes (number of species) per elevation are shown in parentheses.

After correcting for egg mass, GH_2_O differed among stations (Phylogenetic ANOVA; *F*_2,106_ = 29.77, *P* < 0.001), decreasing with increasing elevation. In addition, the variance in GH_2_O differed significantly among stations (*F*_2, 106_ = 4.46, *P* < 0.001), with greater variation in the lowlands and mid-elevations than in the highlands (Fig. 2b). Similar trends (*i*.*e*., greater GH_2_O with greater variability in lowland areas) occurred within families but were not significant; for example, the pattern in the family Tyrannidae resembled the pattern considering species in all families despite being non-significant (Phylogenetic ANOVA; *F*_2,14_ = 3.68, *P* = 0.08; Fig. 2).

### Eggshell structure

Eggshell thickness showed a higher phylogenetic signal (Pagel’s λ = 0.48) than pore density (Pagel’s λ =0.34) or pore size (Pagel’s λ = 0.18), but values λ in all traits were low. This suggests that these eggshell traits are evolutionarily labile (Fig. 3c, f, i), possibly as a result of varying selective pressures among nesting habitats along the mountains.

**Figure 3.**
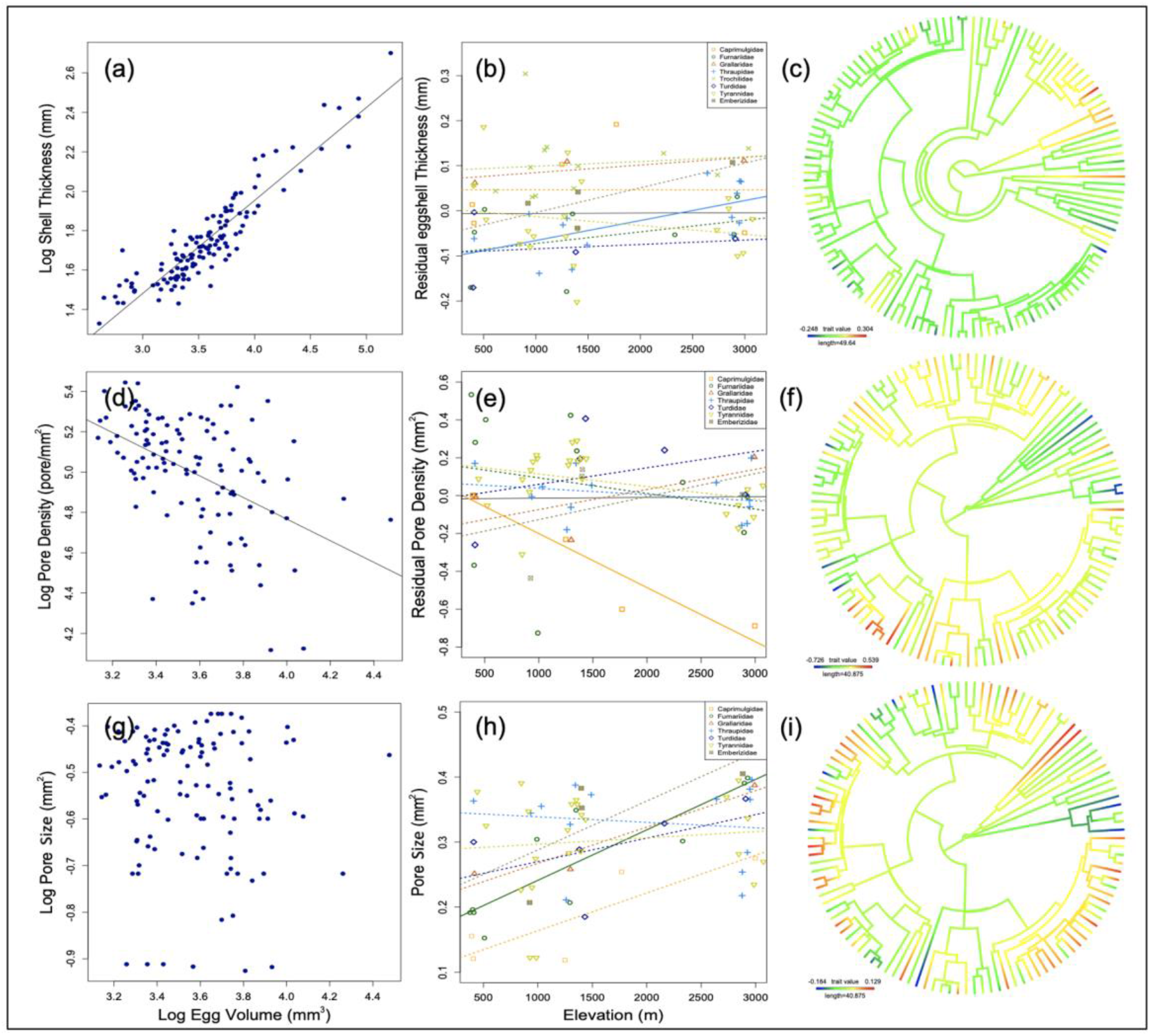
Left: Allometric relationship between (a) thickness, (d) pore density and (g) pore size, and egg volume. Note that thickness is positively correlated with egg size (*P* < 0.001, *n* = 129), while pore density and egg size are negatively related (*P* < 0.001, *n* = 116); there is no relationship between pore size and egg volume, therefore we used absolute values. Center: Relationships between residual (*i*.*e*., size-independent) thickness (b), pore density (e), absolute pore size (h), and elevation for families distributed along the gradient. In all cases, there are no overall relationships, but both positive and negative relationships are found when data are analyzed separately for species in different families: statistically significant relationships are shown as solid lines, otherwise as dotted lines. Right: trait evolution of residual thickness (c), ore density (f), and absolute pore size (i).

Eggshell thickness was positively related to egg volume (*r*^2^ = 0.84, *P* < 0.001; Fig. 3a). PGLS revealed that, across all species, eggshell thickness was not significantly related to elevation after correcting for egg volume (*r*^2^ = 0.007, *P* = 0.053; Fig. 3b). However, eggshell thickness and elevation were positively related among species in Thraupidae (*r*^2^ = 0.33, *P* = 0.018, *n* = 14; Fig. 3b).

Pore density was negatively related to egg volume (*r*^2^ = 0.285, *P* < 0.001; Fig. 3d). Pore density —corrected by egg mass— did not change significantly with elevation across all species (*r*^2^ = 0.01, *P* = 0.99; Fig. 3e). However, there was a negative relationship between residual pore density and elevation in Nightjars (Caprimulgidae; *r*^2^ = 0.87, *P* = 0.014, *n* = 5).

Pore size was not related to egg volume (*r*^2^ = 0.002, *P* = 0.386; Fig. 3g) and was not related to elevation across all species (*r*^2^ =0.004, *P* = 0.51). Within families, we found a positive relationship between pore size and elevation in Furnariidae (*r*^2^ = 0.72, *P* = 0.001, *n* = 10; Fig. 3h).

Within-species analyses did not reveal patterns of variation consistent with adaptation or plasticity in response to elevation. In the Black-faced Brushfinch (*Atlapetes melanolaemus*), we found no significant relationships between elevation and thickness of the shell (*r*^2^ = 0.004, *P* = 0.33), pore density (*r*^2^ = 0.023, *P* = 0.41), or pore size (*r*^2^ = 0.074, *P* = 0.68) over a 1000 m elevation range. Likewise, there was no significant relationships between elevation and shell thickness (*r*^2^ = 0.04, *P* = 0.20, *n* = 19), pore density (*r*^2^ = 0.07, *P* = 0.6), or pore size (*r*^2^ = 0.06, *P* = 0.6) for the Cinnamon Flycatcher (*Pyrrhomyias cinnamomeus*) over a 1700 m elevation range.

## Discussion

Our analyses based on a large sample of Neotropical bird species occurring in montane areas revealed that water vapor conductances across the eggshell are highly labile and vary along elevational gradients in ways consistent with adaptation to better retain humidity in environments with higher partial barometric pressure (Board, 1982; Birchard & Deeming, 2009; Mortola, 2009). A similar conclusion was reached in a study examining global patterns in gas exchange rates across avian eggshells, which documented lower conductances in species breeding at high latitudes, where preventing dehydration is key owing to temperature fluctuations that may suspend embryo growth and prolong incubation (Attard & Portugal, 2021). The results of both comparative studies highlight a prominent role of environmental factors driving adaptations or plasticity in eggshell function across a wide diversity of birds.

Birds from high elevations are expected to compensate for the effects of low ambient water vapor pressure by reducing the eggshell conductance (Rahn et al., 1977; Sotherland et al., 1980; Carey, 1980*a*; Carey et al., 1983; Rahn et al., 1982; Carey et al., 1989). Accordingly, we found that water vapor conductance does decrease with elevation. This pattern occurred among all species and among species within some families but was especially strong in flycatchers and allies (Tyrannidae, Onychorhynchidae). In addition, we found that the variation in conductance was considerably larger at lower elevations. The relatively low variance in conductance at high elevations may reflect environmental filtering, such that only species having particular traits (*i*.*e*. thick shells, low pore density, small pores) may successfully breed at high elevations (Graham et al., 2014).

Alternatively, the higher variation in conductance in lowland and mid-elevation species may reflect relaxed selection and the greater diversity of microclimates (related to greater habitat complexity) at these elevations (Terborgh, 1977; Patterson et al., 1998). Avian diversity in our study region is also highest at low elevations (Terborgh, 1977; Jankowski et al., 2012), and lowland species exhibit a wide spectrum of life-history traits (*e*.*g*., nest construction, nest location, incubation behavior, embryo development period) that may account for the wide variation in water vapor conductance (Deeming & Ferguson, 1991; Deeming, 2002; Mortola, 2009; Attard & Portugal 2021). We acknowledge, however, that because we were unable to keep the same temperature across elevations in experiments at our field sites, decreased conductance in eggs from higher elevations partly reflects an effect of decreasing temperatures with elevation. Although the magnitude of such an effect is unknown, definitive conclusions about variation in mean values of eggshell conductance with elevation thus await additional experiments in which temperature is closely controlled (*see* Carey et al., 1983; Portugal et al., 2010). Nonetheless, the overall pattern of greater variation in conductance we observed in lowland and mid-elevation species cannot be accounted for by variation in temperature across experiments.

Although eggshell conductance decreased with elevation, the mechanisms related to eggshell characteristics potentially underlying this pattern varied among taxa, resulting in no overall relationship between elevation and eggshell characteristics. Birds may achieve reduced GH_2_O at higher elevations via different combinations of thicker eggshells, lower pore density, or reduced pore size. We found two basic patterns in eggshell characteristics consistent with our hypothesis. In tanagers (Thraupidae), shell thickness increased with increasing elevation, whereas pore density and area were constant across elevations. In contrast, in nightjars (Caprimulgidae) and flycatchers and allies (Tyrannidae, Onychorhynchidae) shell thickness decreased slightly with increasing elevation, whereas pore density declined; pore size tended to increase, but not significantly. Differences among families in patterns of variation in eggshell structure along elevational gradients may reflect alternative adaptations to reduce the desiccation (Rahn et al., 1977; Birchard & Deeming, 2009). Presumably adaptive variation, however, was not general. For example, in contrast to predictions, antpittas (Grallariidae), *Turdus* thrushes (Turdidae), and finches (Passerellidae) tended to exhibit higher pore density and area at higher elevations (although patterns were not significant), whereas pore size increased with elevation in ovenbirds (Furnariidae). Given that eggshell structure has likely evolved to meet a variety of different demands (Carey et al., 1983; Birchard & Deeming, 2009), it is noteworthy that we found significant elevational patterns in eggshell function and structure. The significant pattern of changes in eggshell conductance in flycatchers and allies (Tyrannidae, Onychorhynchidae) despite no significant changes in the eggshell structure variables we measured suggests that even minor changes in the eggshell may have important functional consequences for water vapor flux.

Our within-species analyses revealed some variation at the species level in egg structure in species that occupy a wide elevation range. However, within-species variation was unrelated to elevation. This suggests that variation in eggshell structure within populations may allow species to occur over a broad range of elevations, but it is unclear whether the lack of variation with respect to elevation (hence presumably lack of adaptation at range margins) may contribute to setting elevational range limits (Kirkpatrick & Barton, 1997; Stiles, 2004). Possibly other egg traits (*e*.*g*. size) may affect gas exchange across the eggshell and may vary adaptively with elevation. For instance, it has been hypothesized that egg size is larger in species nesting in cool environments to reduce heat loss (Martin, 2008; Heming & Marini, 2015), or that egg mass may increase in the highlands because investing proportionally more water in eggs at laying at higher elevations may compensate for greater water loss during incubation (Carey et al., 1983). Common-garden experiments would be required to directly test for within-species adaptation in egg traits to environmental variables changing with elevation.

### Patterns in eggshell conductance and other selective pressures

Besides pore structure and shell thickness, many other factors may be involved in maintaining the balance between gas diffusion and water flux through the eggshell, and these may differ among taxa (Romanoff & Romanoff, 1949; Becking, 1975). Such factors include incubation behavior (*i*.*e*. nest attendance) and its consequences on incubation temperatures (Carey, 1980b; Carey et al., 1983; Lourens et al., 2005) and the microclimate at nests (Rahn et al., 1982; Portugal et al., 2014). Although we did not find any relationship between nest type and GH_2_O, features such as nest materials used to build nests, nest location and patterns in parental care (Hansell, 2000; Heenan et al., 2015; Gómez *et al*. 2019; Batisteli et al., 2020; Attard & Portugal, 2021) should be taken into account to better describe how nest architecture affects microclimate inside the nest. For example, ovenbirds (Furnariidae) and flycatchers (Tyrannidae), two families showing no changes in eggshell structure with elevation (except for a marginal increase in thickness in flycatchers), vary widely in nest architecture and location (Irestedt et al., 2006; Hemming et al., 2013). Such variation may allow species to adjust conditions to allow for proper embryo development in the absence of changes in the eggshell structure (Collias & Collias, 1984; Hansell, 2000). Another important factor may be the time that the eggs are exposed to the environment, as one would expect that species may reduce exposure times where higher desiccation rates are expected; however, has been suggested that incubation periods tend to be longer at higher elevations in tropical montane birds (Boyle et al., 2015).

The lack of consistent patterns across taxa we observed may also relate to different selective pressures related to abiotic factors driving the evolution of the eggshell in opposite directions. In the temperate zone, low-elevation populations of House Finches (*Haemorhous mexicanus*) exhibit larger eggs with thicker shells and lower pore density relative to populations from higher elevations, which results in lower conductance in the lowlands (Stein & Badyaev, 2011). Given our hypothesis, we expected to find these eggshell characteristics in birds from the highlands, not the lowlands. However, in House Finches a likely more important selective pressure is the risk of trans-shell bacterial infection, which is greater under higher humidity conditions (Cook et al., 2005).

Another possibility is that eggshell structure may not always respond to elevation according to our predictions because of the humidity inside the nest, one of the main drivers of variation in the eggshell structure (Walsberg, 1983; Carey et al., 1983; Portugal et al., 2014; D’Alba et al., 2016) does not necessarily vary linearly with elevation. For instance, species could regulate humidity with their nest architecture, location, and materials (Deeming, 2011), or behaviorally (e.g. incubation patterns), but information on climatic variation at the microscale (i.e. inside the nest) is lacking in our study sites. Future work should pay close attention to the microclimate of individual nests to better characterize the environmental pressures of temperature and water vapor pressure to which eggs are exposed.

### Methodological challenges

A potential caveat of our study is our limited understanding of the functional importance of eggshell structures as quantified via SEM images. For instance, we cannot be sure that all the pores that we detected on the surface penetrated the entire shell. In addition, our basic understanding of the pore structure across the eggshell comes principally from model species, such as chickens and birds in other non-passerine families (Board et al., 1977; Rahn et al., 1979; Tullett, 1984; Vieco-Galvez et al., 2021). We assumed that dark pores penetrate the eggshell, but we acknowledge this is not always true because substances covering the surface may occlude pores (Board et al., 1977; Martín-Vivaldi et al., 2014). The use of techniques that determine which pores are truly functional (e.g., Hargitai et al., 2010; Jaeckle et al., 2012; Bowers et al. 2015) would be a valuable addition to insights obtained based on microscopy. In addition, we were unable to consider details of the three-dimensional structure of egg pores. The 3D structure may influence gas exchange and water loss, although studies on pore morphology suggest that all species in our eggshell structure analysis exhibit simple unbranched canals (Board et al., 1977; Tullett, 1984; Murphy et al., 2015). Further analyses focusing on small species may allow us to understand the trade-off between structural constraints of the eggshell and pore configuration.

### Functional mechanisms. Avian life-history strategies, and elevational distributions

Most studies of variation in avian life-history strategies across elevational gradients have focused on how abiotic and biotic factors affect variables such as fecundity, incubation behavior, clutch size, embryonic and nestling development, and nesting success across multiple species (Badyaev, 1997; Badyaev & Ghalambor, 2001; Boyce et al., 2015; Boyle et al., 2015). However, broad-scale cross-species analyses may conceal patterns within clades that have overcome the same set of evolutionary challenges in different ways (Boyle et al., 2015). Identifying the environmental tolerances of species is critical to understanding the mechanisms underlying the distributions of species with respect to elevation (Jankowski et al., 2013; Graham et al., 2014), but studies on this topic in birds remain scarce (Londoño et al., 2015; 2017; Freeman, 2016). Our work indicates that exploring alternative functional mechanisms such as those related to reproductive traits may provide new perspectives and insight into the factors limiting avian distributions along elevational gradients. Our findings support the hypothesis that characteristics of eggs may have evolved or adjusted via plasticity to match gas exchange rates with local environmental conditions, and that different species may be responding to common pressures in different ways (Board et al., 1977; Board, 1980; Lourens et al., 2005).

Finally, we suggest that examining the extent to which putative adaptations in eggshell function and structure (or the lack thereof) might account for the limits of elevational ranges merits further study. Specifically, providing sufficient oxygen for embryonic development while avoiding desiccation may constrain the distributions of birds at higher elevations and may influence the responses of species to climatic change (Sheldon et al., 2011; Freeman et al., 2018). Given the narrow elevational distribution of species in the tropics and their presumably narrow physiological tolerances (Janzen, 1967), montane species may be especially susceptible to changes in environmental conditions (Pearson & Dawson, 2003; Colwell et al., 2008). Even minor environmental changes may affect microclimates where birds breed and may have deleterious effects on the embryo development (Lourens et al., 2005; Pottier et al. 2022) and species persistence if populations cannot evolve fast enough (or adjust via phenotypic plasticity in structures or behaviors) to cope with changes. This would be especially important in highland areas, where gas conductance through the eggshell shows limited variation. Although other factors (habitat structure, interspecific interactions) have rightly been the focus of most studies on elevational range limits, our work suggests that environmental factors affecting avian eggs also merit consideration.

## Acknowledgments

We thank all the assistants from the Manu Bird Project who helped us in the field in Colombia and Peru, especially Laura Gómez, Andres Chinome, Mario Loaiza, Tim Forrester, and Camilo Florez; it is been emotional! In Peru the assistance of Marianne van Vlaardingen and the Pantiacolla Lodge, Daniel Blanco and the Cock-of-the-Rock Lodge, and the Wayqecha Cloud Forest Biological Station was invaluable. We thank SERNAP for permission to work in the Manu Park buffer zone (0239-2013 MINAGRI-DGFFS/DGEFFS 2013). In Colombia Michelle Tapasco and Gustavo Campuzano helped us with logistics. Scott Whittaker provided technical assistance at the Smithsonian Natural History Museum (NMNH) SEM Lab. Carla Dove and Lorian Straker at the Feather Lab provided very helpful guidance and comments during the process of studying eggshell structures. Jacob Saucier helped us during the data collection at the egg collection at the NMNH. Esteban Correa and Marcela Hernández made valuable contributions to developing the algorithm that we used for the data collecting of the eggshell pore characteristics. Simón Quintero helped us collect data on eggshell thickness. Liam Revell and Luke Harmon helped with the comparative analysis throughout Latin Amer. Macroevolution Workshop 2014. We thank members of the “Laboratorio de Biología Evolutiva de Vertebrados” for very productive discussions. Mary C. Stoddard, David Outumuro, Mark Chappell, Robert Ricklefs, Scott Robinson, Juan A. Amat, Tony D. Williams, Maria C. Estrada-F, Zuania Colon, Juliana Soto, Orlando Acevedo, and anonymous reviewers provided valuable comments on the manuscript. Our animal use was approved by the University of Florida IACUC and ACCA. Funding was provided by “Proyecto Semilla” from the Facultad de Ciencias at Universidad de Los Andes, the Francois Vuilleumier Fund from the Neotropical Ornithological Society, The American Ornithologists’ Union Research Award, Graduate Student Fellowships – Smithsonian Institute, and the National Science Foundation Grant DEB-1120682.

## Authors’ contributions

All the authors conceived the ideas and designed the methodology.

David O and Gustavo L collected the data.

All the authors analysed the data.

David O led the writing of the manuscript.

All authors contributed critically to the drafts and gave final approval for publication.

## Data availability statement

Data for this paper will be deposited in the Dryad Digital Repository.

## Literature Cited

Ar, A., Paganelli, C. V., Reeves, R. B., Greene, D. G. & Rahn. H. (1974) The avian egg: water vapor conductance, shell thickness, and functional pore area. Condor, 76, 153–158.

Attard, M. R., & Portugal, S. J. (2021) Climate variability and parent nesting strategies influence gas exchange across avian eggshells. Proceedings of the Royal Society B, 288(1953), 20210823.

Attard, M. R., Bowen, J., Corado, R., Hall, L. S., Dorey, R. A., & Portugal, S. J. (2021) Ecological drivers of eggshell wettability in birds. Journal of the Royal Society Interface, 18(183), 20210488.

Attard, M. R., & Portugal, S. J. (2022) Global diversity and adaptations of avian eggshell thickness indices. Ibis. doi.org/10.1111/ibi.13136

Badyaev, A. V. (1997) Covariation between life history and sexually selected traits: an example with cardueline finches. Oikos, 80, 128–138.

Badyaev, A. V. & Ghalambor, C. K. (2001) Evolution of life histories along elevational gradients: trade-off between parental care and fecundity. Ecology, 82,2948–2960.

Batisteli, A. F., de Souza, L. B., Santieff, I. Z., Gomes, G., Soares, T. P., Pini, M., … & Sarmento, H. (2020) Buildings promote higher incubation temperatures and reduce nest attentiveness in a Neotropical thrush. Ibis. 163(1), 79–89.

Becking, J. H. (1975) The ultrastructure of the avian eggshell. Ibis, 117,143–151.

Birchard, G. F. & Deeming, D. C. (2009) Avian eggshell thickness: scaling and maximum body mass in birds. Journal of Zoology 279, 95–101.

Board, R. G. (1980) The avian eggshell—a resistance network. Journal of Applied Bacteriology, 48, 303–313.

Board, R. G., & Scott, V. D. (1980) Porosity of the avian eggshell. American Zoologist, 20(2), 339–349.

Board, R. G. (1982) Properties of avian eggshells and their adaptive value. Biological Reviews, 57, 1–28.

Board, G. R., Tullett, G. S. & Perrott. R. H. (1977) An arbitrary classification of the pore systems in avian eggshells. Journal of Zoology, 182, 251–265.

Bowers, E. K., White, A., Lang, A., Podgorski, L., Thompson, C. F., Sakaluk, S. K., … & Harper, R. G. (2015) Eggshell porosity covaries with egg size among female House Wrens (Troglodytes aedon) but is unrelated to incubation onset and egg-laying order within clutches. Canadian journal of zoology, 93(6), 421–425.

Bota-Sierra, C. A., García-Robledo, C., Escobar, F., Novelo-Gutiérrez, R., & Londoño, G. A. (2022) Environment, taxonomy, and morphology constrain insect thermal physiology along tropical mountains. Functional Ecology, 36(8), 1924–1935.

Boyce, A. J., Freeman, B. G., Mitchell, A. E. & Martin, T. E. (2015) Clutch size declines with elevation in tropical birds. The Auk, 132, 424–432.

Boyle, A. W., Sandercock, B. K. & Martin, K. (2015) Patterns and drivers of intraspecific variation in avian life history along elevational gradients: a meta-analysis. Biological Reviews, 000–000.

Butler, M. A. & King, A. A. (2004) Phylogenetic Comparative Analysis: A Modeling Approach for Adaptive Evolution. The American Naturalist, 164, 683–695.

Cadena, C. D. & Loiselle, B. A. (2007) Limits to elevational distributions in two species of emberizine finches: disentangling the role of interspecific competition, autoecology, and geographic variation in the environment. Ecography, 30, 491–504.

Carey, C. (1980a) Adaptation of the avian egg to high altitude. American Zoologist, 20, 449–459.

Carey, C. (1980b) The Ecology of Avian Incubation. BioScience, 30, 819–824.

Carey, C., Leon-Velarde, F. & Monge, C. (1990) Eggshell conductance and other physical characteristics of avian eggs laid in the Peruvian Andes. Condor, 92, 790–793.

Carey, C., Leon-Velarde, F. & Dunin-Borkowski, O. (1989) Variation in eggshell characteristics and gas exchange of montane and lowland coot eggs. Journal of Comparative Physiology B, 159, 389–400.

Carey, C., Thompson, E. L., Vleck, C. M. & James, F. C. (1982) Avian reproduction over an altitudinal gradient: incubation period, hatchling mass, and embryonic oxygen consumption. The Auk, 99, 710–718.

Carey, C., Garber, S. D., Thompson, E. L. & James, F. C. (1983) Avian reproduction over an altitudinal gradient. II. Physical characteristics and water loss of eggs. Physiological zoology, 56, 340–352.

Collias, N. E., & Collias, E. C. (1984) Nest building and bird behavior. Princeton University Press.

Colwell, R. K., Brehm, G., Cardelus, C. L., Gilman, A. C. & Longino, J. T. (2008) Global Warming, Elevational Range Shifts, and Lowland Biotic Attrition in the Wet Tropics. Science, 322, 258–261.

Cook, M. I., Beissinger, S. R., Toranzos, G. A., Rodriguez, R. A. & Arendt, W. J. (2005) Microbial infection affects egg viability and incubation behavior in a tropical passerine. Behavioral Ecology, 16, 30–36.

D’Alba, L., Maia, R., Hauber, M. E., & Shawkey, M. D. (2016) The evolution of eggshell cuticle in relation to nesting ecology. Proc. R. Soc. B, 283, 1836, 20160687.

Deeming, D. C. (2002) Avian incubation: behaviour, environment and evolution. Oxford: Oxford University Press.

Deeming, D. C. (2011) Importance of nest type on the regulation of humidity in bird nests. Avian Biology Research, 4(1), 23–31.

Deeming, D. C., & Ferguson., M. W. J. (1991) Egg Incubation. Cambridge University Press.

Diamond, J. M. (1973) Distributional ecology of New Guinea birds. Science, 179, 759–769.

Duursma, D. E., Gallagher, R. V., Price, J. J., & Griffith, S. C. (2018) Variation in avian egg shape and nest structure is explained by climatic conditions. Scientific reports, 8(1), 4141.

Fecheyr-Lippens, D. C., Igic, B., D’Alba, L., Hanley, D., Verdes, A., Holford, M., Waterhouse G., Grim T., Hauber M., & Shawkey, M. D. (2015) The cuticle modulates ultraviolet reflectance of avian eggshells. Biol Open 4 (7), 753–759. doi.org/10.1242/bio.012211.

Felsenstein, J. (1973) Maximum-likelihood estimation of evolutionary trees from continuous characters. American journal of human genetics, 25, 471–492.

Fierro-Calderón, K., Loaiza-Muñoz, M., Sánchez-Martínez, M. A., Ocampo, D., David, S., Greeney, H. F., & Londoño, G. A. (2021) Methods for collecting data about the breeding biology of Neotropical birds. Journal of Field Ornithology. doi.org/10.1111/jofo.12383

Freckleton, R. P., Harvey, P. H. & Pagel, M. (2002) Phylogenetic Analysis and Comparative Data: A Test and Review of Evidence. The American Naturalist, 160, 712–726.

Freeman, B. G. (2016) Thermal tolerances to cold do not predict upper elevational limits in New Guinean montane birds. Diversity and Distributions, 22,309–317.

Freeman, B. G., Lee-Yaw, J. A., Sunday, J. M., & Hargreaves, A. L. (2018) Expanding, shifting and shrinking: The impact of global warming on species’ elevational distributions. Global Ecology and Biogeography, 27(11), 1268–1276.

Freeman, B. G., Tobias, J. A., & Schluter, D. (2019) Behavior influences range limits and patterns of coexistence across an elevational gradient in tropical birds. Ecography, 42(11), 1832–1840.

Freeman, Benjamin G., Matthew Strimas-Mackey, and Eliot T. Miller. (2022) Interspecific competition limits bird species’ ranges in tropical mountains. Science 377.6604, 416–420.

Funk, W. C., & Biellier, H. V. (1944) The minimum temperature for embryonic development in the domestic fowl (Gallus domesticus). Poultry science, 23, 538–540.

García-Robledo, C., Kuprewicz, E. K., Staines, C. L., Erwin, T. L. & Kress, W. J. (2016) Limited tolerance by insects to high temperatures across tropical elevational gradients and the implications of global warming for extinction. Proceedings of the National Academy of Sciences of the USA, 113, 680–685.

Ghalambor, C. K., Huey, R. B., Martin, P. R., Tewksbury, J. J & Wang, G. (2006) Are mountain passes higher in the tropics? janzen’s hypothesis revisited. Integrative and Comparative Biology, 46, 5–17.

Graham, C. H., Carnaval, A. C., Cadena, C. D., Zamudio, K. R., Roberts, T. E., Parra, J. L., McCain, C. M. et al. (2014) The origin and maintenance of montane diversity: integrating evolutionary and ecological processes. Ecography, 37, 711–719.

Gómez, J., Liñán-Cembrano, G., Ramo, C., Castro, M., Pérez-Hurtado, A., & Amat, J. A. (2019) Does the use of nest materials in a ground-nesting bird result from a compromise between the risk of egg overheating and camouflage?. Biology open, 8(12).

Hackett, S. J., Kimball, R. T., Reddy, S., Bowie, R. C. K., Braun, E. L., Braun, M. J., Chojnowski, J. L. et al. (2008) A Phylogenomic Study of Birds Reveals Their Evolutionary History. Science, 320, 1763–1768.

Hansell, M. (2000) Bird Nests and Construction Behaviour. Cambridge University Press.

Hargitai, R., Mateo, R. & J. Török. (2010) Shell thickness and pore density in relation to shell colouration, female characteristics, and environmental factors in the Collared Flycatcher Ficedula albicollis. Journal of Ornithology, 152, 579–588.

Harmon, L. J., J. B. Losos, T. Jonathan Davies, R. G. Gillespie, J. L. Gittleman, W. Bryan Jennings, K. H. Kozak, et al. (2010) Early bursts of body size and shape evolution are rare in comparative data. Evolution, 64, 2385–2396.

Harmon, L. J., Weir, J. T., Brock, C. D., Glor, R. E. & Challenger, W. (2007) GEIGER: investigating evolutionary radiations. Bioinformatics, 24, 129–131.

Heenan, C. B., Goodman, B. A., & White, C. R. (2015) The influence of climate on avian nest construction across large geographical gradients. Global Ecology and Biogeography, 24, 11, 1203–1211.

Heming, N. M. & Marini, M. Â. (2015) Ecological and environmental factors related to variation in egg size of New World flycatchers. Journal of Avian Biology, 46, 001–009.

Irestedt, M., Fjeldså, J. & Ericson, P. G. (2006) Evolution of the ovenbird-woodcreeper assemblage (Aves: Furnariidae)-major shifts in nest architecture and adaptive radiation. Journal of Avian Biology, 37, 260–272.

Jaeckle, W. B., Kiefer, M., Childs, B., Harper, R. G., Rivers, J. W., & Peer, B. D. (2012) Comparison of eggshell porosity and estimated gas flux between the brown-headed cowbird and two common hosts. Journal of Avian Biology, 43(6), 486–490.

Jankowski, J., Merkord, C. L., Rios, W. F., Cabrera, K. G., Revilla, N. S. & Silman, M. R. (2012) The relationship of tropical bird communities to tree species composition and vegetation structure along an Andean elevational gradient. Journal of Biogeography, 40, 950–962.

Jankowski, J. E., Londoño, G. A., Robinson, S. K. & Chappell, M. A. (2013) Exploring the role of physiology and biotic interactions in determining elevational ranges of tropical animals. Ecography, 36, 1–12.

Janzen, D. H. (1967) Why mountain passes are higher in the tropics. The American Naturalist, 101, 233–249.

Jetz, W., Thomas, G. H., Joy, J. B., Hartmann, K. & Mooers, A. O. (2012) The global diversity of birds in space and time. Nature, 491, 444–448.

Karlsson, O., & Lilja, C. (2008) Eggshell structure, mode of development and growth rate in birds. Zoology, 111, 494–502.

Kirkpatrick, M. & Barton, N. H. (1997) Evolution of a species’ range. The American Naturalist, 150, 1–23.

Linck, E. B., Freeman, B. G., Cadena, C. D., & Ghalambor, C. K. (2021) Evolutionary conservatism will limit responses to climate change in the tropics. Biology Letters, 17(10), 20210363.

Londoño, G. A., Chappell, M. A., Castañeda, M. D. R., Jankowski, J. E. & Robinson, S. K. (2015) Basal metabolism in tropical birds: latitude, altitude, and the “pace of life.” Functional Ecology, 29, 338–346.

Londoño G.A., Chappell M.A., Jankowski J.E., & Robinson S.K. (2017) Do thermoregulatory costs limit altitude distributions of Andean forest birds? Functional Ecology, 31, 204–2015.

Lourens, A., Van den Brand, H., Meijerhof, R. & Kemp, B. (2005) Effect of eggshell temperature during incubation on embryo development, hatchability, and posthatch development. Poultry science, 84, 914–920.

Martin, T. E. (2008) Egg size variation among tropical and temperate songbirds: an embryonic temperature hypothesis. Proceedings of the National Academy of Sciences, 105, 9268–9271.

Martín-Vivaldi, M., Soler, J. J., Peralta-Sánchez, J. M., Arco, L., Martín-Platero, A. M., Martínez-Bueno, M., Ruiz-Rodriguez, M. et al. (2014) Special structures of hoopoe eggshells enhance the adhesion of symbiont-carrying uropygial secretion that increase hatching success. Journal of Animal Ecology, 83, 1289–1301.

Maurer, G., Portugal, S. J., Mikšík, I. & Cassey, P. (2011) Speckles of cryptic black-headed gull eggs show no mechanical or conductance structural function. Journal of Zoology, 285, 194–204.

McCracken, K. G., Bulgarella, M., Johnson, K. P., Kuhner, M. K., Trucco, J., Valqui, T. H., Wilson, R. E. et al. (2009) Gene Flow in the Face of Countervailing Selection: Adaptation to High-Altitude Hypoxia in the A Hemoglobin Subunit of Yellow-Billed Pintails in the Andes. Molecular Biology and Evolution, 26, 815–827.

Mikhailo, K. (1997) Avian Eggshells: an Atlas of Scanning Electron Micrographs. British Ornithologists’ Club, 3, 87 pg.

Mortola, J. P. (2009) Gas exchange in avian embryos and hatchlings. Comparative Biochemistry and Physiology, Part A, 153, 359–377.

Murphy, J. P., Swanson, M. T., Jaeckle, W. B. & Harper, R. G. (2015) Corrosion casts: A novel application of a polyurethane resin (PU4ii) for visualizing eggshell pore morphology. The Auk 132, 206–211.

Narushin, V. G. (2005) Egg geometry calculation using the measurements of length and breadth. Poultry science, 84, 482–484.

Orme, D. (2013) The caper package: comparative analysis of phylogenetics and evolution in R. R package version, 5, 1–36.

Österström, O., & Lilja, C. (2011) Evolution of avian eggshell structure. Journal of Morphology, 273, 241–247.

Paganelli, C. V., Ar, A. & Lanphieh E. H. (1971) The influence of pressure and gas composition on water vapor diffusion. Proc. Int. Union Physiol. Sci, 9, 1294.

Paganelli, C. V., Ar, A., & Rahn, H. (1973) Relative water vapor permeabilities of the shell and shell membranes of the hen’s egg. Physiologist, 16, 415.

Paganelli, C. V., Olszowka, A. & Ar, A. (1974) The avian egg: surface area, volume, and density. Condor, 76, 319–325.

Pagel, M. (1999) Inferring the historical patterns of biological evolution. Nature, 401, 877–884.

Patterson, B. D., Stotz, D. F., Solari, S., Fitzpatrick, J. W. & Pacheco, V. (1998) Contrasting patterns of elevational zonation for birds and mammals in the Andes of southeastern Peru. Journal of Biogeography, 25, 593–607.

Pearson, R. G. & Dawson, T. P. (2003) Predicting the impacts of climate change on the distribution of species: are bioclimate envelope models useful?. Global ecology and biogeography, 12, 361–371.

Portugal, S. J., Maurer, G. & Cassey, P. (2010) Eggshell Permeability: A Standard Technique for Determining Interspecific Rates of Water Vapor Conductance. Physiological and Biochemical Zoology, 83, 1023–1031.

Portugal, S. J., Maurer, G., Thomas, G. H., Hauber, M. E., Grim, T. & Cassey, P. (2014) Nesting behaviour influences species-specific gas exchange across avian eggshells. Journal of Experimental Biology, 217, 3326–3332.

Pottier, P., Burke, S., Zhang, R. Y., Noble, D. W., Schwanz, L. E., Drobniak, S. M., & Nakagawa, S. (2022) Developmental plasticity in thermal tolerance: Ontogenetic variation, persistence, and future directions. Ecology Letters, 25, 2245–2268.

Rahn, H., Ar, A. & Paganelli, C. V. (1979) How bird eggs breathe. Scientific American, 240, 46–55.

Rahn, H., Carey, C., Balmas, K., Bhatia, B. & Paganelli, C. (1977) Reduction of pore area of the avian eggshell as an adaptation to altitude. Proceedings of the National Academy of Sciences of the USA, 74, 3095–3098.

Rahn, H. & Ar, A. (1980) Gas Exchange of the Avian Egg Time, Structure, and Function. Integrative and Comparative Biology, 20, 477–484.

Rahn, H., Ledoux, T., Paganelli, C. V. & Smith, A. H. (1982) Changes in eggshell conductance after transfer of hens from an altitude of 3,800 to 1,200 m. Journal of applied physiology: respiratory, environmental and exercise physiology, 53, 1429–1431.

Rahn, H. & Paganelli, C. V. (1990) Gas fluxes in avian eggs: driving forces and the pathway for exchange. Comparative Biochemistry and Physiology Part A: Physiology, 95, 1–15.

Rasband, W. S. (2008) ImageJ. http://rsbwebnihgov/ij/.

Revell, L. J. (2010) Phylogenetic signal and linear regression on species data. Methods in Ecology and Evolution, 1, 319–329.

Revell, L. J. (2012) phytools: an R package for phylogenetic comparative biology (and other things). Methods in Ecology and Evolution, 3, 217–223.

Rokitka, M.A. & Rahn, H. (1987) Regional differences in shell conductance and pore density of avian eggs. Respir. Physiol. 68, 371–376. doi:10.1016/S0034-5687(87)80021-X

Romanoff, A. L. (1930) Biochemistry and biophysics of the developing hens’ egg. I. Influence of humidity. Agricultural Experimental Station Memoirs, 132, 1–27 (Cornell University, New York State College of Agriculture).

Romanoff, A. L., & Romanoff, A. J. (1949) The avian egg. John Wiley & Son, New York, NY.

Sheldon, K. S., Yang, S. & Tewksbury, J. J. (2011) Climate change and community disassembly: impacts of warming on tropical and temperate montane community structure. Ecology Letters, 14, 1191–1200.

Slatyer, R. A. & Schoville, S. D. (2016) Physiological Limits along an Elevational Gradient in a Radiation of Montane Ground Beetles. PLoS ONE, 11, e0151959.

Sotherland, P. R., Packard, G. C., Taigen, T. L. & Boardman, T. J. (1980) An altitudinal cline in conductance of cliff swallow (Petrochelidon pyrrhonota) eggs to water vapor. The Auk, 97, 177–185.

Stein, L. R. & Badyaev, A. V. (2011) Evolution of eggshell structure during rapid range expansion in a passerine bird. Functional Ecology, 25, 1215–1222.

Stiles, F. G. (2004) Phylogenetic constraints upon morphological and ecological adaptation in hummingbirds (Trochilidae): Why are there no hermits in the paramo. Ornitologia Neotropical, 15, 191–198.

Stoddard, M. C., Yong, E. H., Akkaynak, D., Sheard, C., Tobias, J. A., & Mahadevan, L. (2017) Avian egg shape: Form, function, and evolution. Science, 356(6344), 1249–1254.

Storz, J. F., Scott, G. R. & Cheviron, Z. A. (2010) Phenotypic plasticity and genetic adaptation to high-altitude hypoxia in vertebrates. Journal of Experimental Biology, 213, 4125–4136.

Terborgh, J. (1971) Distribution on environmental gradients: theory and a preliminary interpretation of distributional patterns in the avifauna of the Cordillera Vilcabamba, Peru. Ecology, 52, 23–40.

Terborgh, J. (1977) Bird species diversity on an Andean elevational gradient. Ecology, 58, 1007–1019.

Tullett, S.G. (1984) The porosity of avian eggshells. Comparative Biochemistry and Physiology Part A: Physiology, 78, 5–13.

Tullett, S. G. & Board, R. G. (1977) Determinants of avian eggshell porosity. Journal of Zoology, 183, 203–211.

Vieco-Galvez, D., Castro, I., Morel, P. C., Chua, W. H., & Loh, M. (2021) The eggshell structure in apteryx; form, function, and adaptation. Ecology and Evolution, 11(7), 3184–3202.

Vleck, C. E. M., Hoyt D.F. & Vleck D. (1979) Metabolism of avian embryos: patterns in altricial and precocial birds. Physiol. Zool., 52, 363–377.

Vleck, C. M., Vleck, D., Rahn, H. & Paganelli, C. V. (1983) Nest microclimate, water-vapor conductance, and water loss in heron and tern eggs. The Auk, 100, 76–83.

Walsberg, G. E. (1983) A test for regulation of nest humidity in two bird species. Physiological zoology, 56, 231–235.

Wangensteen, O. D., Wilson, D. & Rahn, H. (1971) Diffusion of gases across the shell of the hen’s egg. Respiration physiology, 11, 16–30.

Wangensteen, O. D., Rahn, H., Burton, R. R., & Smith, A. H. (1974) Respiratory gas exchange of high altitude adapted chick embryos. Respiration physiology, 21(1), 61–70.

White, C.R., Blackburn, T.M., Martin, G.R. & Butler, P.J. (2007) Basal metabolic rate of birds is associated with habitat temperature and precipitation and not primary productivity. Proceedings of the Royal Society of London Series B, 274, 287–293.

Zuloaga, J., & Kerr, J. T. (2016) Over the top: do thermal barriers along elevation gradients limit biotic similarity?. Ecography, 40(4), 478–486.

